# Human–wildlife conflict shows shared spatial predictors across ecological and geographic contexts

**DOI:** 10.64898/2026.10.10.758226

**Authors:** Tanner C. Smith, Enrico Di Minin, Naomi Kodama, Nicolas Ray, Takuya Iwamura

## Abstract

Human-wildlife conflict (HWC) is a global challenge for biodiversity conservation, food security, and human well-being. Effective management towards human-wildlife coexistence, recognized as a priority in Target 4 of the Kunming–Montreal Global Biodiversity Framework, requires understanding the drivers of HWC. However, the strong influence of local human and ecological context means that most HWC research focuses on individual case studies, limiting transferability of patterns across contexts. Here, we link local contexts and transferable drivers by testing whether HWC exhibits an intermediate level of organization, in which local predictor profiles are structured by broader ecological, geographic, and taxonomic contexts. After compiling ∼11,000 conflict incidents, we construct and compare random forest models across 29 mammalian HWC scenarios spanning diverse species, conflict types, biomes, and continents to identify contexts in which predictor profiles converge. While predictors varied among individual conflicts, HWC scenarios were organized by shared context: conflicts occurring within the same biome tended to have similar predictor profiles, with continent and taxonomic group providing additional structure. Proximity to protected areas was the most consistently important predictor of HWC globally. The likelihood of conflict increased with human population density across nearly all scales and contexts, while distance to roads was also an important predictor in many contexts. Together, these results demonstrate an intermediate level of organization in HWC that reconciles case-specific variation with broader patterns across biomes, continents, and taxonomic groups. This framework provides a basis for scaling mitigation strategies beyond individual case studies while avoiding one-size-fits-all approaches.

## Introduction

Human–wildlife conflict (HWC) is a major challenge for sustainable development, impacting both biodiversity conservation and human well-being goals^1,2^. Crop and livestock losses, threats to livelihoods, and retaliatory killing of wildlife can undermine coexistence and erode support for conservation^2–5^. Reconciling the dissonance between global conservation goals and the lived experience of those that interact with wildlife the most makes monitoring and predicting HWC to mitigate conflict paramount for achieving the sustainable development goal. Understanding the locations and predictors of HWC enables governments and communities to allocate funding and resources to mitigate the worst effects of conflict, however monitoring is often insufficient on its own^6^. Predictive models enable preventative management and land-use planning^6–8^. This challenge is increasingly relevant to global conservation policy, including the Kunming–Montreal Global Biodiversity Framework, which explicitly recognizes human–wildlife coexistence within its biodiversity targets^9,10^.

HWC occurs within diverse social-ecological systems, and most research has focused on individual case studies tailored to local conditions^11^. Local case studies yield important insights, but due to the inherent complexity of HWC, their application outside of their immediate context is limited^12^. Comparisons are further complicated by differences in data sources, sampling methods, spatial scales, and the covariates considered^13^. Although recent global reviews have synthesized HWC research^14^, the diversity of approaches and disciplines makes it difficult to determine which ecological and spatial predictors consistently characterize conflict across contexts^15^. Here, we focus specifically on spatial and ecological predictors of conflict, providing a common framework for evaluating their relevance across otherwise disparate HWC studies.

Potential predictors of HWC include human population density and agricultural activity, wildlife habitat and resources, protected areas, terrain, and infrastructure, but their relevance varies among species, landscapes, and study systems^16–20^. This variation raises a central question of this study: which features of HWC are context-specific, and which recur across systems? Increasing availability of HWC data and globally standardized environmental covariates now makes it possible to address this question across a broader range of contexts^15,21^. Following a paradigm developed in Sociology and recently applied in sustainable sciences that aims to bridge empirical observation with theory, we hypothesized that HWC exhibits an intermediate level of organization^22,23^, in which individual conflicts remain strongly shaped by local conditions but predictor profiles are shared among conflicts occurring within similar ecological, geographic, or taxonomic contexts. We therefore use a meta-analytical framework to test whether biome, continent, taxonomic group, and conflict type explain similarities in predictor profiles across HWC studies (Figure 1).

**Figure 1.**
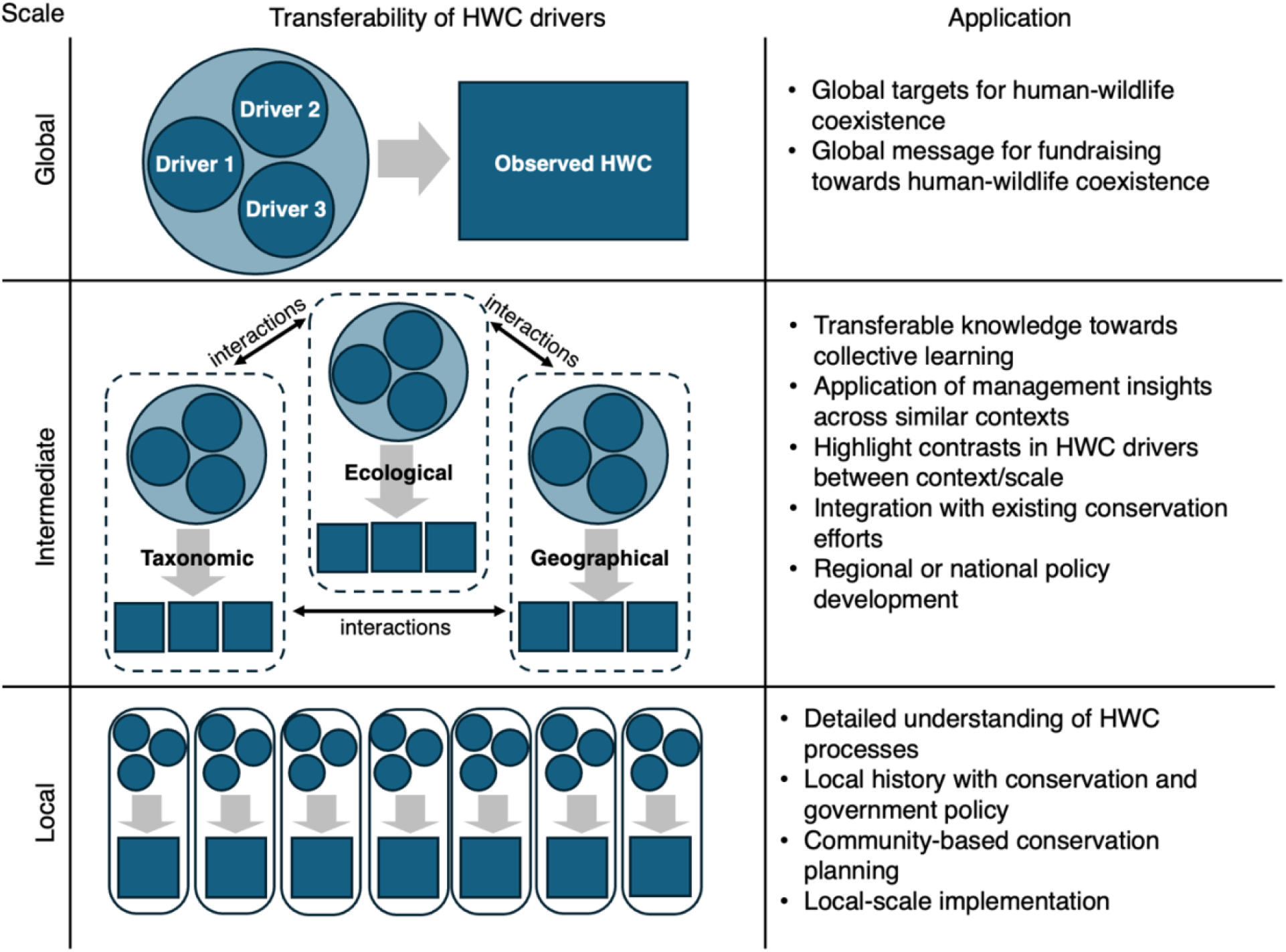
Conceptual diagram of intermediate interpretive framework of HWC. We assume that the predictors of HWC may vary at different spatial and contextual scales. Circles represent predictors of HWC, and squares represent HWC. Unique context can be expressed via multiple predictors in statistical models. Our work demonstrates an intermediate scale where shared variables can account for multiple realities when grouped in appropriate contexts such as biome, taxa, or continent.

To determine which of the contexts share spatial predictors of HWC and which predictors are broadly relevant across systems, we combined a global database of mammalian HWC with standardized spatial modeling. Building on our systematic literature review^14^, we compiled more than 11,000 conflict incidents from 29 unique contexts worldwide^24–44^, and developed probabilistic random forest models for each scenario using nine globally standardized predictors. We then examined whether similarities in predictor profiles were structured by biome, continent, taxonomic group, or conflict type. This framework allows us to distinguish predictors that recur across diverse HWC contexts while identifying the intermediate ecological and geographic scales at which local patterns may be similar. Rather than treating HWC as either a collection of isolated case studies or a globally uniform phenomenon, we use this approach to test how local variation is organized within broader patterns of HWC.

## Results

Our literature search yielded 20 papers with available data matching our criteria (Table 1)^24–44^, containing data for 29 unique HWC contexts, when divided by taxa and conflict types. The datasets were available for 12 countries, namely India, Italy, Romania, Poland, Thailand, Nepal, China, United States of America, Canada, Mozambique, Botswana, and Zimbabwe. We created models for each of the conflict scenarios in each of the study areas to predict HWC probability based on nine key predictive variables which recur in HWC modeling literature (Figure 2), and which also minimized the variable inflation factor of the models to ensure the predictive variables were not redundant. We found there to be high variability in the accuracy of the models as assessed by model performance (AUC-PR) ranging from 0.56-0.9. Accuracy of the models was correlated with sample size (Figure S1).

**Table 1.** Studies compiled for spatially explicit HWC dataset

| Publication | Periods | HWC types | HWC species | Biome | Country | N |
| --- | --- | --- | --- | --- | --- | --- |
| Athreya et al. 2015<sup>25</sup> | 2013-2014 | Livestock predation | Leopard | Tropical forest | India | 151 |
| Baral et al. 2021<sup>26</sup> | 2011-2019 | Human safety | Asian black bear | Subtropical forest | Nepal | 76 |
| Bautista et al. 2021<sup>27</sup> | 2014-2017 | Livestock predation | Brown bear | Temperate forest | Romania | 123 |
| Branco et al. 2021<sup>28</sup> | 2016-2017 | Crop damage | African elephant | Savannah and dry scrub | Mozambique | 1662 |
| Chaudhuri et al. 2022<sup>29</sup> | 2021 | Human safety | Sloth bear | Subtropical forest | India | 56 |
| Davoli et al. 2022<sup>30</sup> | 2009-2021 | Crop damage, livestock predation | Wild boar, Gray wolf | Temperate forest | Italy | 3507 |
| Farr et al. 2022<sup>31</sup> | 2010-2020 | Nuisance | Coyote | Cold coniferous forest | Canada | 726 |
| Karki & Panthi 2021<sup>32</sup> | 2020 | Livestock predation | Snow leopard | Cold coniferous | Nepal | 109 |
| Loveridge et al. 2019<sup>33</sup> | 2009-2016 | Livestock predation | Lion, Spotted hyena, Cheetah, Caracal, Leopard | Savannah and dry scrub | Zimbabwe | 2929 |
| Malpeli et al. 2020<sup>34</sup> | 2008-2015 | Crop damage, Nuisance, Livestock predation, Property damage | American black bear | Temperate forest | USA | 4880 |
| Malviya & Kirishnamurthy 2022<sup>35</sup> | 2021 | Livestock predation | Tiger | Tropical forest | India | 110 |
| Mekonnen et al. 2012<sup>36</sup> | 2012 | Crop damage | Vervet monkey | Subtropical forest | Ethiopia | 36 |
| Naha et al. 2021<sup>37</sup> | 2015-2019 | Crop damage | Asian elephant | Subtropical forest | India | 508 |
| Pop et al. 2023<sup>38</sup> | 2008-2016 | Crop damage | Brown bear | Temperate forest | Poland | 750 |
| Simon & Fortin 2020<sup>39</sup> | 2013-2016 | Crop damage | American bison | Cold coniferous forest | India | 154 |
| Treves & Rabenhorst 2017<sup>40,41</sup> | 2003-2009 | Livestock predation | Grey wolf | Temperate forest | USA | 167 |
| Vasudev et al. 2022<sup>42</sup> | 2013-2021 | Property damage, Human safety, Crop damage | Asian elephant | Tropical forest | India | 1482 |
| Weise et al. 2020<sup>43</sup> | 2016-2018 | Livestock predation | Lion | Savannah and dry scrub | Botswana | 196 |
| Wettasin et al. 2023<sup>44</sup> | 2000-2020 | Crop damage | Asian elephant | Tropical forest | India | 460 |
| Yang et al. 2024 | 2023 | Crop damage | Wild boar | Subtropical forest | China | 638 |

**Figure 2.**
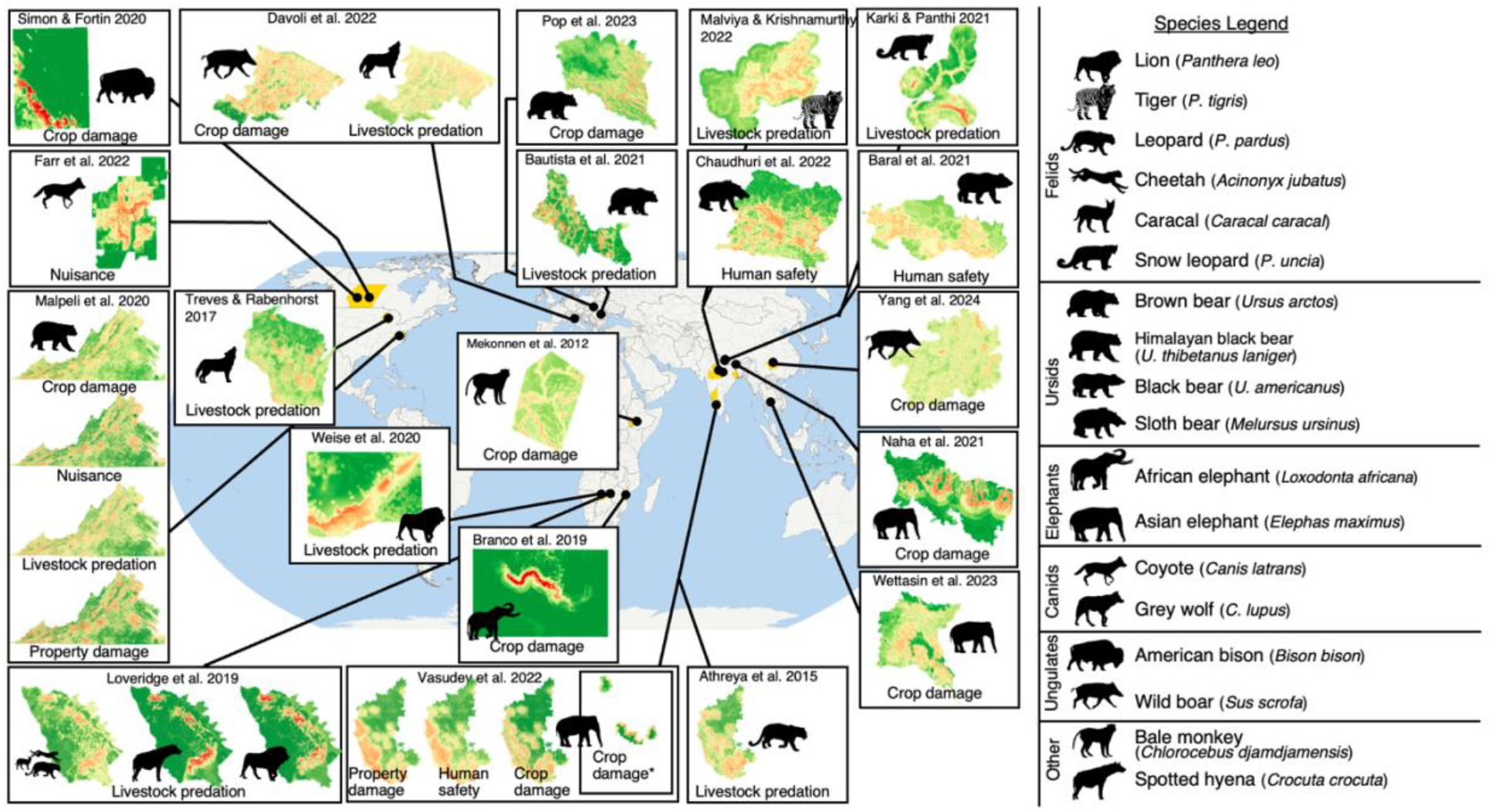
Global map of predicted conflict risk using meta spatial analysis. Each box represents a distinct study. Based on the reported HWC occurrence from these studies are modelled using predictors at local scale (protected area distance, distance from roads, population density, for the full list see methods). Each map corresponds to a specific conflict scenario of unique combination of species, location, or conflict type. Within each predictive map occurrence for HWC, red represents high conflict risk and green represents areas of low conflict risk. Dark silhouette represents the focal species for each study, detailed on right side.

The importance of all predictive variables varied by individual study, but patterns formed depending on groupings within contextual variables (Figure 3, 4). We found protected area distance, human population density, and road distance to be the three most consistently influential predictive variables across contexts globally. The importance of predictive variables in models from Asia demonstrated high variability, but the three most influential were consistent here, along with terrain ruggedness and water distance (Figure 3d). Meanwhile, the models for Africa were most influenced by protected area distance, road distance, and distance to agriculture. The models for Europe and North America resembled one another, with both having tree canopy cover, and population density as the two other most consistently influential predictive variables (Figure 4a).

**Figure 3.**
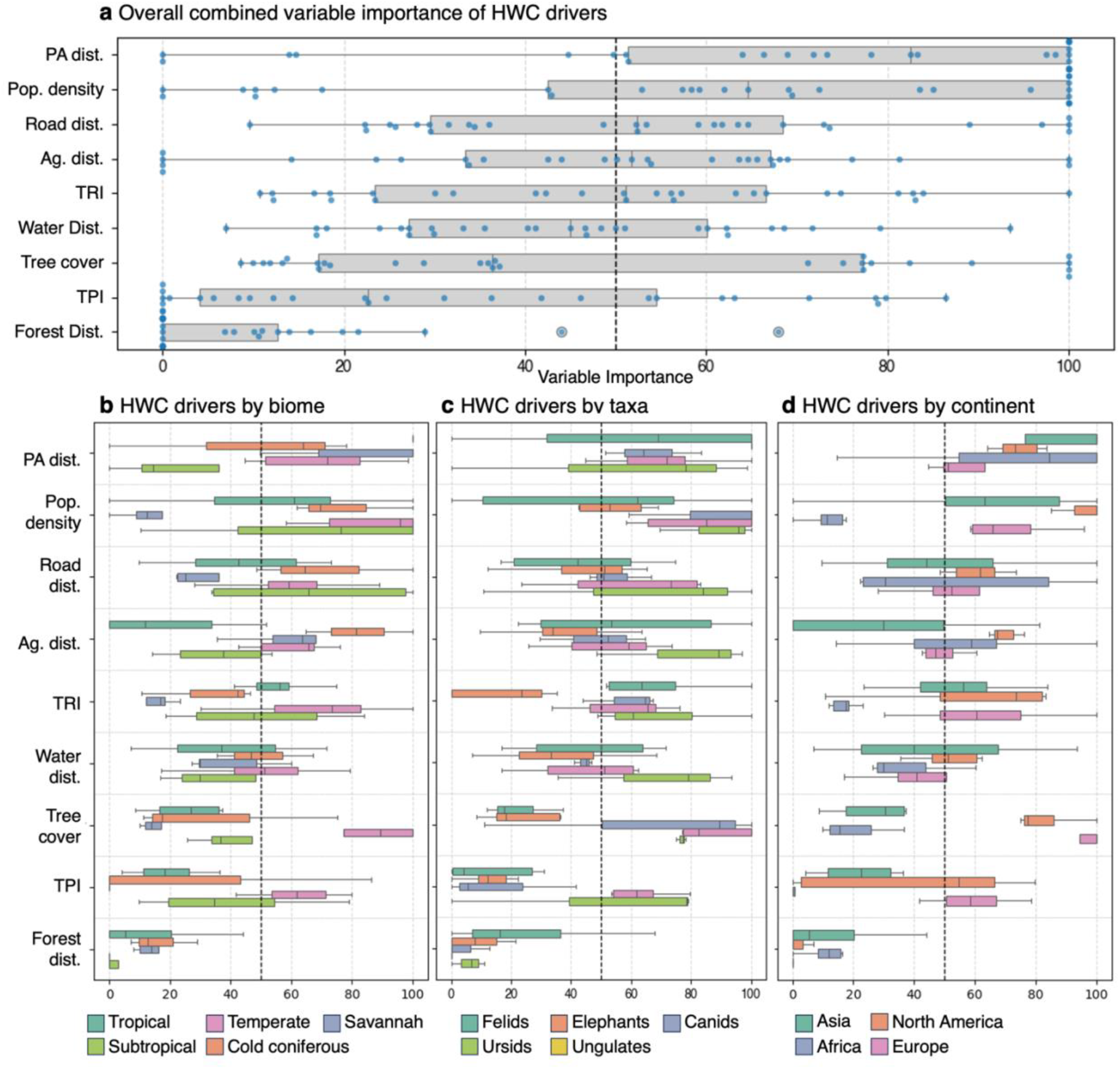
Boxplots describing predictive variable importance across models. In all charts the normalized importance (0-100) is represented on the x-axis. The predictive variables appear on the y-axis in order of median importance across all models. a. Model importances overall and uncategorized, each point represents a model. b. grouped boxplots of simplified biome groupings. c. grouped boxplots of taxonomic groupings. d. grouped boxplots of continental groupings

**Figure 4.**
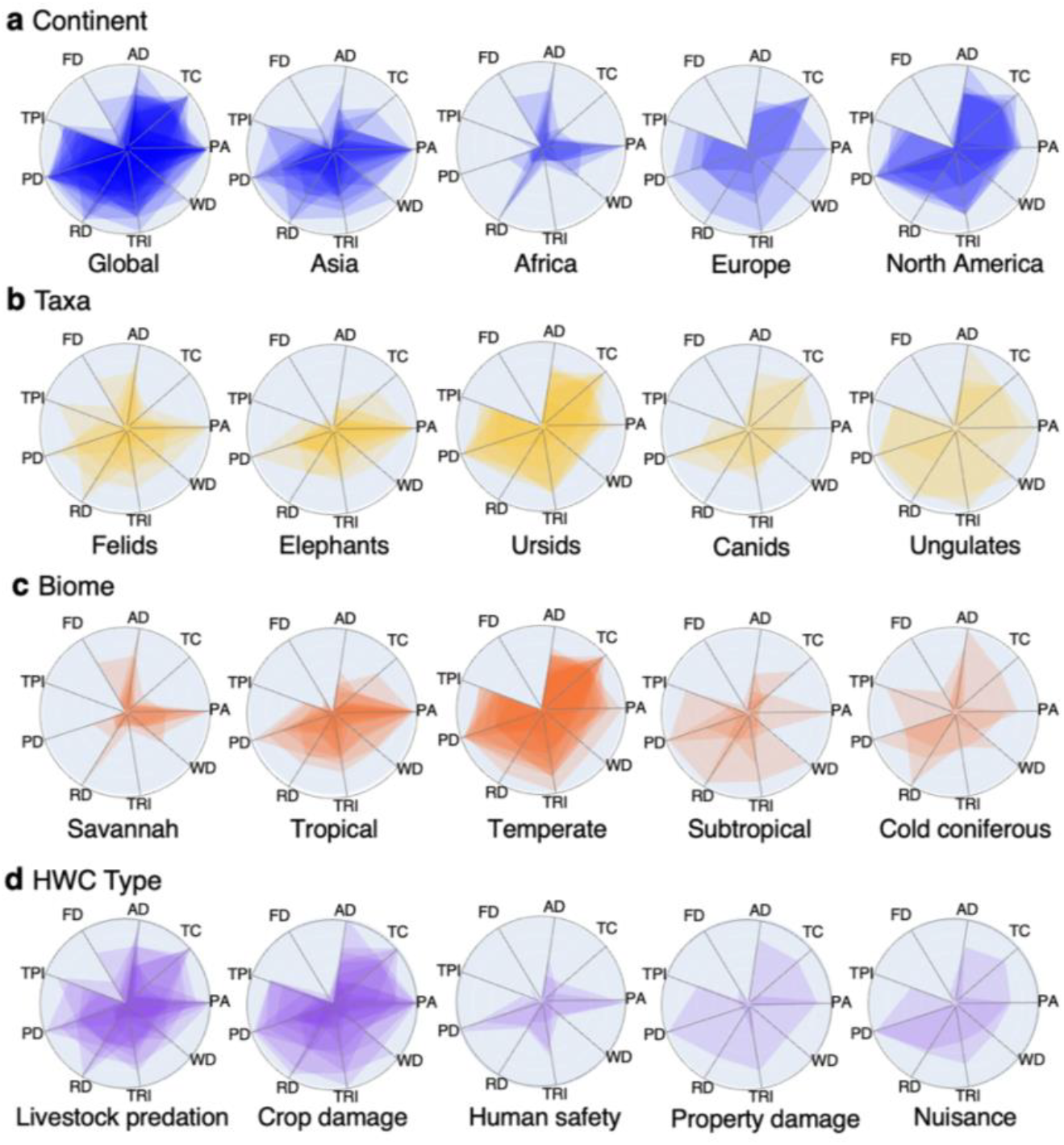
Radar charts of variable importance grouped by intermediate scale variables. Radar plots demonstrate the normalized value (0-100) of variable importance for each of the nine local variables used for each model. The shape of each model within a category is overlapped, opacity indicates variables with frequent agreement for that category. Variable values are noted as the following: FD - forest distance, AD - agricultural distance, TC - tree canopy cover, PA - protected area distance, WD - distance to Water, TRI - terrain ruggedness index, RD - distance to road, PD - human population density, TPI - topographic position index. a. Plots grouped by continent, with all models included in the global category. b-d. Radar charts grouped by taxa, biome, and HWC type respectively.

The variable importance in the biome groupings exhibited the greatest variation for variable importance as the variables are most consistently influential globally were not as consistent across biomes (Figure 3b). For example, here PA distance varies between groups and in subtropical forests it is one of the least important predictive variables. Conversely, tree cover a variable that is not important overall is one of the most important variables in temperate forest biomes, along with population density (Figure 4c).

Across taxonomic groups, distance to protected areas and human population density were the most consistently important correlates of conflict (Figure 3c). Felids were the primary exception, with four different predictors ranking highest across five models, indicating substantially greater variability than other taxa. In contrast, elephant, ursid, canid, and ungulate models showed more consistent predictor profiles across species and locations. Their strongest distinguishing correlates included terrain ruggedness for elephants, tree canopy cover and distance to agricultural land for ursids, and tree canopy cover for canids and ungulates (Figure 4b).

Four unsupervised clusters formed when we compared the influence of predictive variables amongst our models, in most cases conflict scenarios from the same study area were the most closely related. Although geographical proximity plays a role, data collection methods also play a role despite some variance in these groupings as well (Figure 5). Notably, regardless of taxonomic group or conflict type, the national income level also played a role in the similarity of the models. European and North American models were clustered together whereas models of conflict in Africa and India were more similar in terms of predictive variable influence.

**Figure 5.**
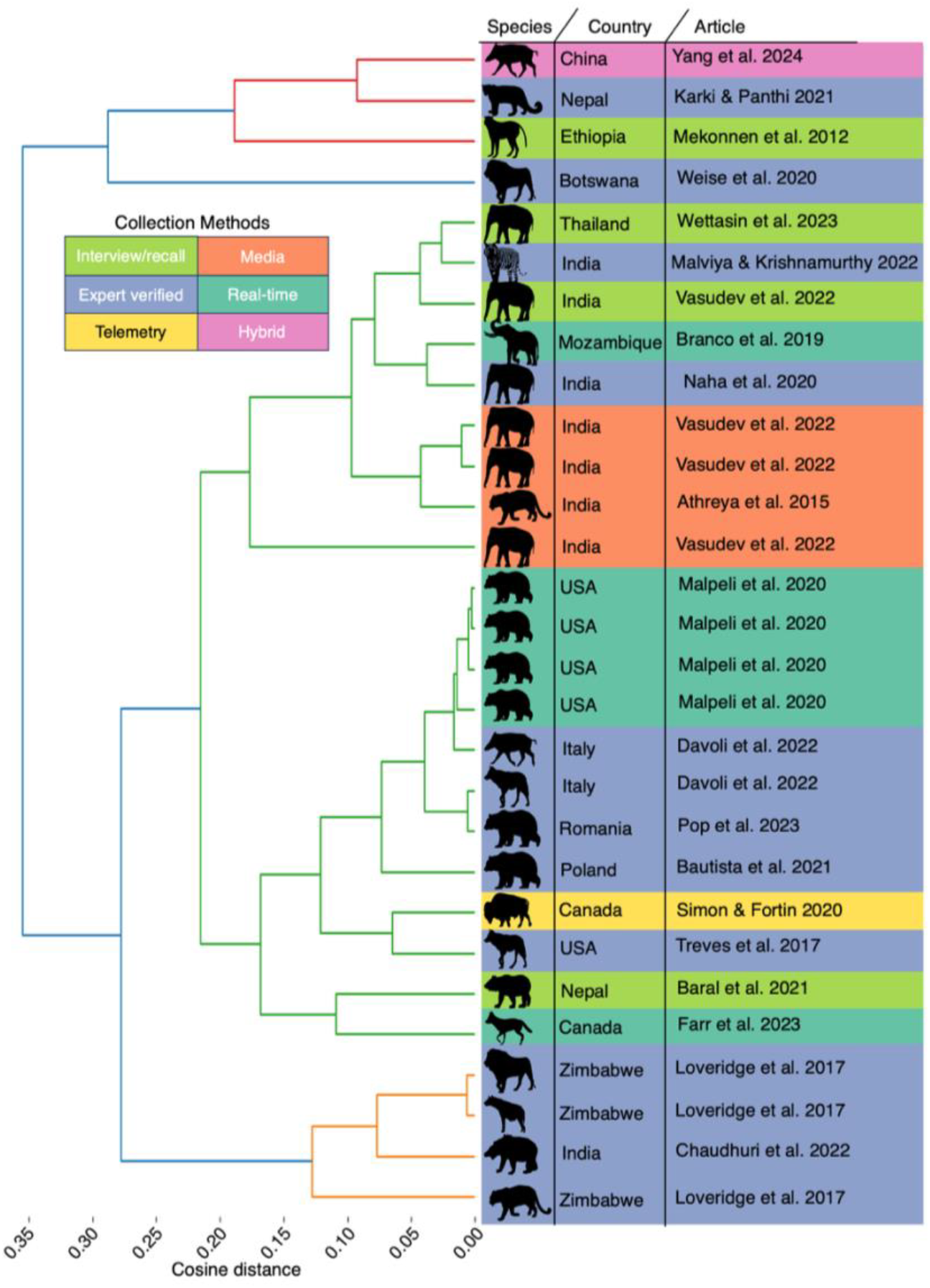
Dendrogram of clustered models based on predictive variable importance. Species involved in conflict and the location of the modelled area are all reflected in this chart. Species silhouettes are defined in figure 2. Branch colors represent 4 distinct clusters based upon cosine distances and average linkage clustering derived from Random Forest model predictive variable importances. Colors on the right side of the figure represent the collection method of the data collection from the original dataset (interviews, media sources, expert monitoring or report verification, real time reporting by community members, telemetry or proxy-based measurements, hybrid combinations of approaches).

The differences in the models were largely driven by the variability of the utility of protected area distance, population density, topographic components, and road distance in model performance (Figure 4). One of the largest clusters included the models with relatively low protected area influence and high influence of terrain ruggedness index, population density, tree canopy cover, and topographic position index. Although not universally impactful, agricultural and forest distance both drive variance in some cases, particularly Africa-based scenarios (Figure 5a).

Our PERMANOVA test found that out of 4 contextual variables hypothesized to explain the clustering (biome, continent, taxa, HWC type), all except HWC type explained the variance in the data through group centroid divergence (Figure 6). The biome in which the conflict was documented stood out as the most clearly distinct group centroids. This is notable due to the geographical breadth of the biomes, as biomes are not exclusive to a particular region or continent. Felid species were amongst the least predictable taxa, as the models defining conflict in the group are highly context dependent. In many instances, outliers within these hypothesized clusters can be explained by another contextual variable, demonstrating their interconnectedness.

**Figure 6.**
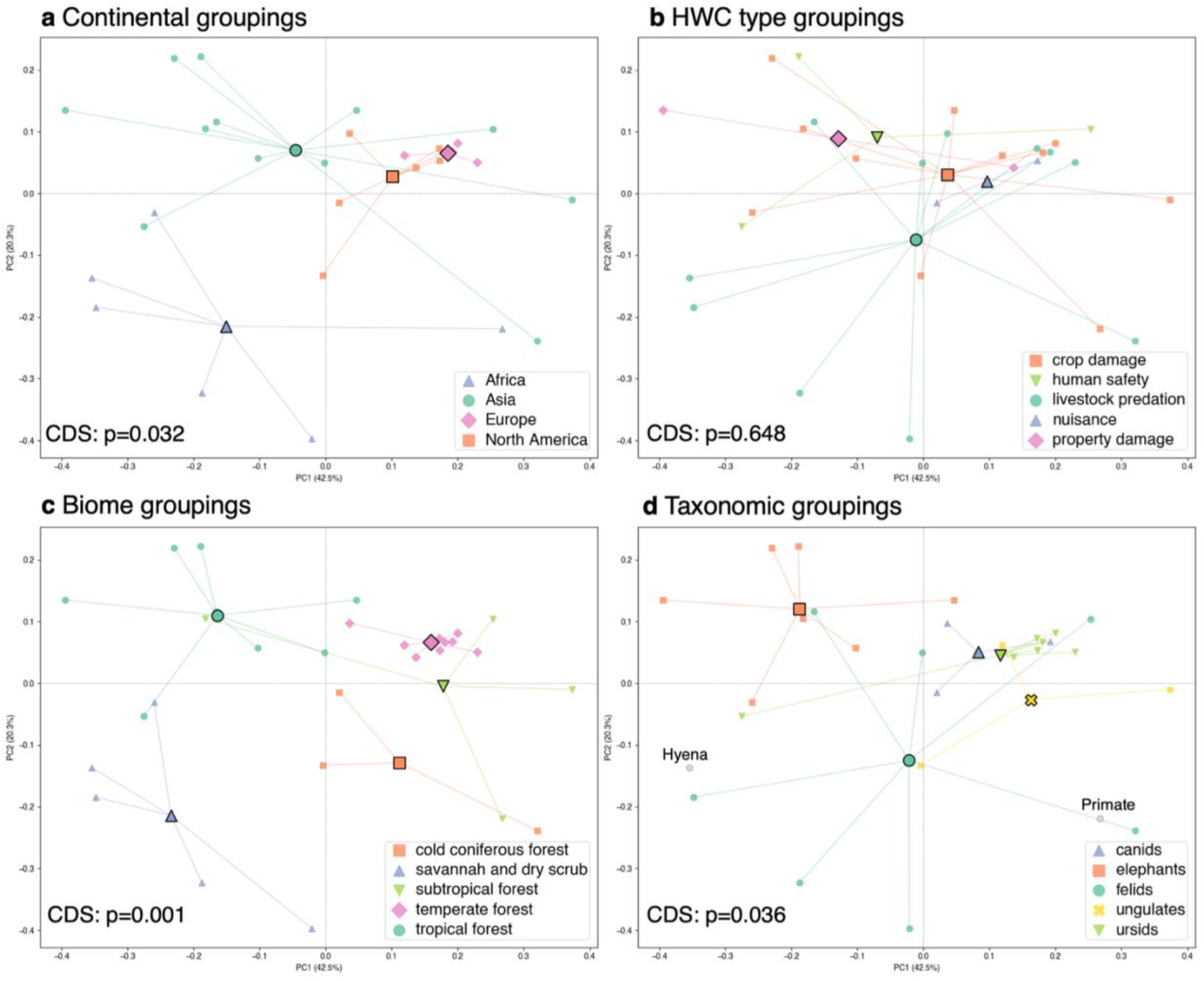
Principal coordinate analysis biplots clustered at intermediate scales. These plots demonstrate the variance of the first two dimensions of multidimensional space for model comparisons. Percentages indicate variance explained on each axis. Intermediate scale variable groupings represented in spider clusters for continent, conflict type, biome, and species groupings respectively. Lines indicate relationship of individual studies to group centroids. PERMANOVA centroid difference significance (CDS) for each variable grouping is represented in the bottom left of each chart, a value of p<0.05 indicates significant difference between group centroid values. a. Studies grouped by continent. b. Studies grouped by Human-wildlife conflict type. c. Studies grouped by biome. d. Studies grouped by taxonomic group. Uncategorized hyenas and primates are not grouped but represented on the plot.

## Discussion

Our results reveal an intermediate level of organization in the predictive models of human–wildlife conflict (HWC), which means although individual conflicts remain strongly shaped by local context, their predictor profiles exhibit broader structure across biomes, continents, and taxonomic groups (Figure 1). Across diverse contexts, spatial patterns of HWC were associated with a small set of globally standardized predictors, with models produced from the same study area sharing the most predictors, followed by studies that fall within the same category of the intermediate levels of organization (Figure 3-6). Taxonomic group and continent further explained variation in these patterns. Nevertheless, study area remained the dominant source of variation in most contexts, highlighting that broad-scale regularities do not replace the need to understand local conditions. The identification of an intermediate scale of related HWC patterns open the door to potentially more precisely and robustly applying research findings in multiple contexts.

While our results highlight the importance of intermediate scale organization, the consistency of several predictors across otherwise disparate HWC scenarios points to processes that transcend local or regional conflict contexts. Human population density, distance to protected areas, and distance to roads were among the most consistently important predictors globally (Figure 3, 4), suggesting that human land use and accessibility are common features of HWC across systems. At the same time, the continental separation in predictor profiles—with North American and European models more similar to one another than to those from Asia or Africa—indicates that these common relationships are embedded within broader geographic differences (Figure 6).

Although HWC arises from context-specific social, cultural, and ecological processes^11,20^, our results identify several predictors that recur across otherwise disparate contexts. Human population density and distance to protected areas were among the most consistently important predictors (Figure 3), reflecting the fundamental role of spatial overlap between people and large mammals in creating opportunities for conflict^19,45^. Protected areas may be particularly relevant because many large mammals depend on extensive habitat for foraging, seasonal movements, and dispersal^46–48^. Other indicators of wildlife habitat or presence, including tree cover, forest distance, and water distance, were less consistently important across models, suggesting that their relevance is more context dependent. These findings highlight the importance of anticipating HWC when establishing area-based conservation measures in landscapes shared with human populations^17^.

The global consistency of these predictors was nevertheless structured by intermediate-scale context. Biome, taxonomic group, and continent explained variation in predictor profiles, whereas conflict type did not (Figure 6), indicating that where conflict occurs and which taxa are involved may be more informative than the type of conflict itself. Distance to protected areas was most relevant in Africa and Asia, where many of the large-bodied and threatened species represented in our analyses occur, whereas its relevance was lower in Europe and North America (Figure 3, 4). In these regions, conflicts more often involve generalist species and wildlife populations recovering in human-dominated or mixed landscapes^49–51^. These differences may reflect variation in species assemblages, landscape structure, conservation history, and human land use across regions.

Our findings have direct implications for conservation planning under the Kunming–Montreal Global Biodiversity Framework, which explicitly recognizes human–wildlife coexistence alongside ambitious area-based conservation targets^10^. Because protected areas and human population density were among the most consistent predictors of HWC (Figure 3), potential conflict should be considered explicitly when identifying and expanding protected areas, particularly in landscapes where large rural populations depend on agriculture or livestock^1,18^. At the same time, the strong influence of local context indicates that mitigation and management of HWC are likely to be more effective when tailored to the relevant biome, taxonomic group, and local human and land-use conditions. Integrating HWC predictors into conservation decision-making protocol could therefore help better identify protected-area locations, and also development strategies that reduce conflict while meeting conservation objectives^52–54^.

Several limitations may constrain the generality of the patterns reported here. Our database lacked sufficient data from South and Central America, Central Asia, and parts of Africa, while differences in data availability and sample size limited model development in some regions (Figure S1). The underlying studies may also vary in sampling and reporting methods, while we addressed this as much as possible via sub-municipal thresholds. Expanding standardized HWC monitoring, including through reporting frameworks such as SMART and IUCN guidelines, would improve comparability among studies and allow broader geographic coverage^55,56^. Integrating grey literature and locally published data (e.g., compensatory records) could further reduce geographic and language biases, although such sources require careful assessment of reporting bias^13,57^.

HWC action plans face a policy–research gap, whereby globally coordinated actions must be informed by an understanding of localized case studies. Our results demonstrate that these perspectives are not mutually exclusive, where HWC exhibits broad-scale structure while remaining strongly shaped by local context. Identifying intermediate spatial and conceptual scales provides a framework for reconciling these patterns, allowing global models to inform regional conservation planning while retaining the context needed for effective local mitigation. Recognizing intermediate scales of organization in HWC provides a pathway to bridge local conflict dynamics with global conservation priorities, enabling interventions that are transferable without losing the ecological and social context in which conflicts occur.

## Methods

We created a modeling pipeline to predict spatial probability of the occurrence of human wildlife conflict (Figure S2). We trained a random forest model using existing and published data on human and mammalian wildlife conflict. All covariate data used for prediction is publicly available at the global scale, enabling simple integration of new datasets and study regions into the modeling pipeline. The output of the classification model was then used to identify significant trends and correlates of human-wildlife conflict.

### Study areas

For each dataset included we looked at the literature and attempted to directly extract their study area or georeferenced it as closely as possible to what was represented in the original research (Figure 2). This was a key step for ensuring accuracy of the models and that the conflict data was as representative as possible.

The study areas range in size from 206 km2 to 191,811 km2 and span the globe with Africa, Asia, Europe, and North America represented. The study areas also range from heterogenous wildland and urban areas to sparsely populated rural or wild areas.

### Data acquisition and cleaning

We used a structured query to identify literature on human-wildlife conflict of terrestrial vertebrates as part of a systematic scoping review^14^. We identified additional datasets using a snowballing approach to identify any data not officially published in a full academic article. We screened the literature for articles with published data or data available upon request. Authors were systematically contacted to request data and available data was downloaded from the relevant repositories.

Only datasets containing spatially explicit data at a scale finer than the municipal level was considered for inclusion. Specifically, we retained only incidents involving terrestrial mammals and conflict that could be classified as crop damage, attack on humans, livestock predation, or nuisance wildlife incidents. We cleaned the data to maintain only the species involved, the type of conflict, the timing of the conflict, and the GPS location of the conflict in World Geodetic System 1984 (WGS 84) as well as metadata of the source. We completed all data cleaning using Python 3.12 and Quantum Geographic Information Systems (QGIS 3.40.5). No data was excluded based upon geographic location. We included only datasets with at least 20 separate recorded instances of HWC. Occurrence points were assessed after being filtered so that all points were at least 100 meters apart.

### Covariates

We selected covariates based upon their ecological relevance as well as their occurrence in previous literature surrounding prediction of human-wildlife conflict ^58–68^. We used Google Earth Engine’s Python implementation to gather, and consolidate the datasets used in the model. Most datasets included in our analysis are readily available in Google Earth Engine’s data catalog, as well as the Earth Engine Community Catalog, any datasets downloaded from other public sources were included in GEE as private assets. We incorporated predictive variables relating to land cover and land use by creating Euclidean distance rasters indicating the distance outside a given feature (e.g., distance to nearest road), pixels within a given class are assigned a zero value. All predictive and contextual variables considered and their data sources are found in Table S1.

### Variable selection

For each of the candidate predictive variables, we calculated their variance inflation factor (VIF); by taking 5000 random points throughout the study area, we evaluated the interrelationships of the correlates. VIF is a measure of multicollinearity which examines the relationship of all other predictive variables to the target variable, going beyond pairwise correlation^69^. VIF above 5 indicates that the remaining predictive variables combine to explain a large portion of the variance of the target predictive variable and therefore may distort model interpretation. Although predictive accuracy is unlikely to be affected by collinearity in random forest models, the interpretation of predictive variable importance can be affected^70^. We removed predictive variables that caused VIF to exceed 5 in most cases and 10 in unique cases (Table S2)^71^. We found there was no single set of predictive variables that was able to completely avoid exceeding VIF threshold values in all study areas. We found landscape fragmentation predictive variables to consistently increase the total VIF in most study areas, so we removed them from analysis.

### Pseudo-negative points

We categorized areas as negative that were outside of a 100-meter buffer surrounding our occurrence points, and randomly sampled these areas to produce an equal number of occurrence points and pseudo-negatives to resemble a down-sampled random forest model^72–74^. We used the down-sampled random-forest model to predict the probability of conflict across each study area based upon the correlated variables included in 100-meter pixels^75,76^. We divided each landscape into blocks for spatial block cross validation, to split species occurrence datasets into training and testing datasets of 70%-30% respectively, while accounting for spatial autocorrelation^77,78^. In our multi-model workflow, we found that the performance of the model was highly dependent on the size of the grid, so we performed a sensitivity analysis for each model for hyperparameter tuning to identify which grid size between 5000, 10000, 20000, 30000, 40000, and 50000 meters allowed the model to perform best according to area-under-the-curve precision-recall (AUC-PR) of the output model average after 3 runs at each level.

### Model Comparison

For each area, using the optimized grid size, we trained 10 down-sampled random forest models on random combinations of training and testing spatial blocks, the data ensuring each of the training and testing splits contained at least 40 training and 20 testing points to avoid empty training or testing sets. We averaged the results of the 10 models to obtain an average of all outputs (predictions, AUC-PR, predictive variable importance). We identified variable importance by using permutation importance on each of the predictive variables to identify their influence on the accuracy of the model. The average importance was then rescaled from 0-100 to enable comparison between models of relative variable importance. We then made a distance matrix of all the models, by calculating the cosine distance between each model according to the importance. Next, we completed a principal coordinate analysis on the square root of the distance matrix to identify the drivers of variance within the models and contexts^79^. Finally, we conducted Permutational Multivariate Analysis of Variance (PERMANOVA) test to identify which contextual variables available at the multiple study level had categories significantly different from one another. We chose human-wildlife conflict, taxonomic group (hyenas and primates formed a group called “other”), the majority biome, and the continent of the study area as the contextual variables to understand the multivariate centroid differences of interpretable groupings.

## Acknowledgements

We would like to thank all data providers and field workers for their important work in collecting the data used here, as well as their commitment to open science. In particular, we would like to thank the CLAWS Conservancy as well as the National Geographic Society and Stichting SPOTS, Netherlands.

## Author Contributions

Conceptualization: T.C.S., E.D.M., N.R., T.I.; Methodology: T.C.S., T.I.; Date collection: T.C.S., N.K.; Investigation: T.C.S., E.D.M., T.I.; Programming: T.C.S.; Visualization: T.C.S., T.I.; Funding acquisition and project administration: T.I.; T.C.S. wrote the initial draft and all authors contribute to the writing, review and editing of the original draft.

