## Supplemental Information for "Human–wildlife conflict shows shared spatial predictors across ecological and geographic contexts"

**Supporting Figures**

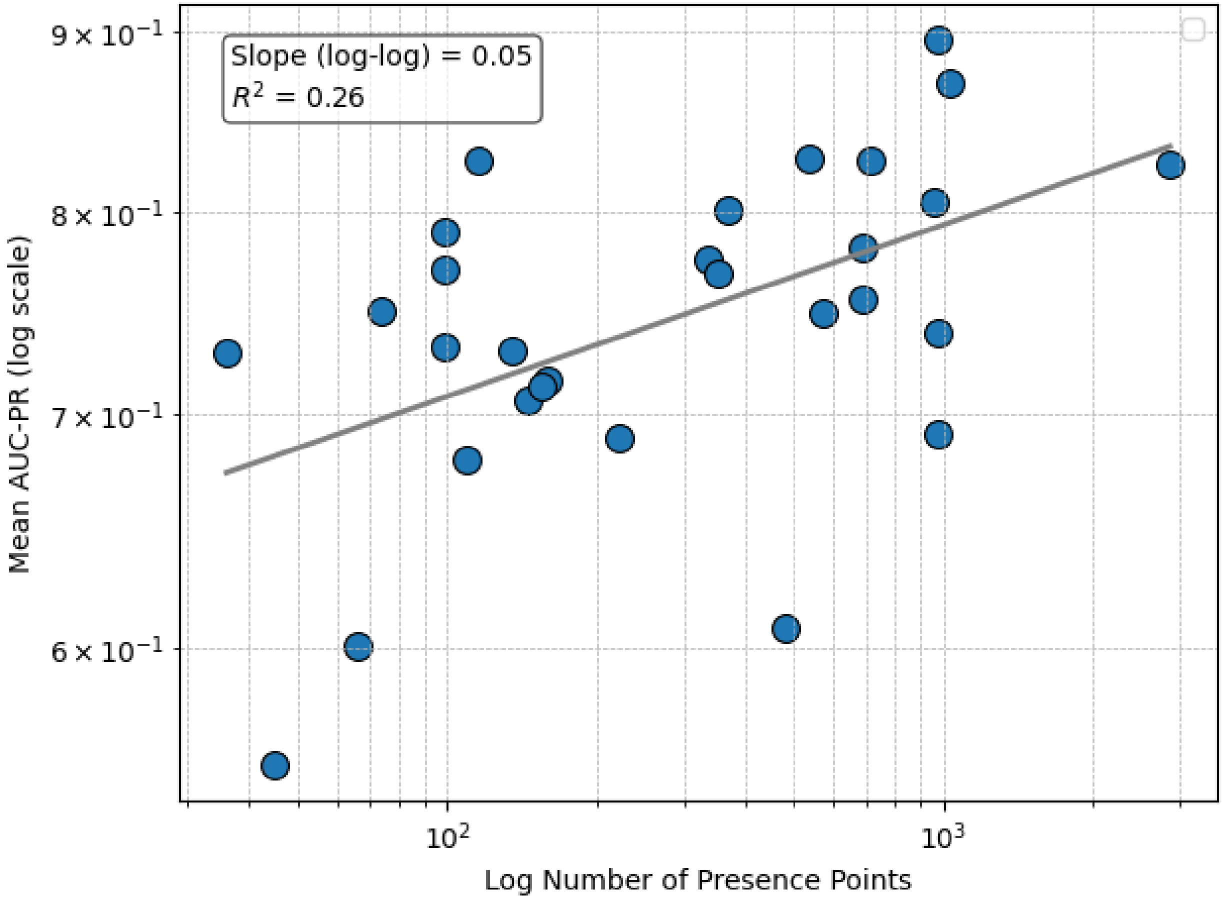


**Figure S1.** Log relationship between model accuracy and sample size

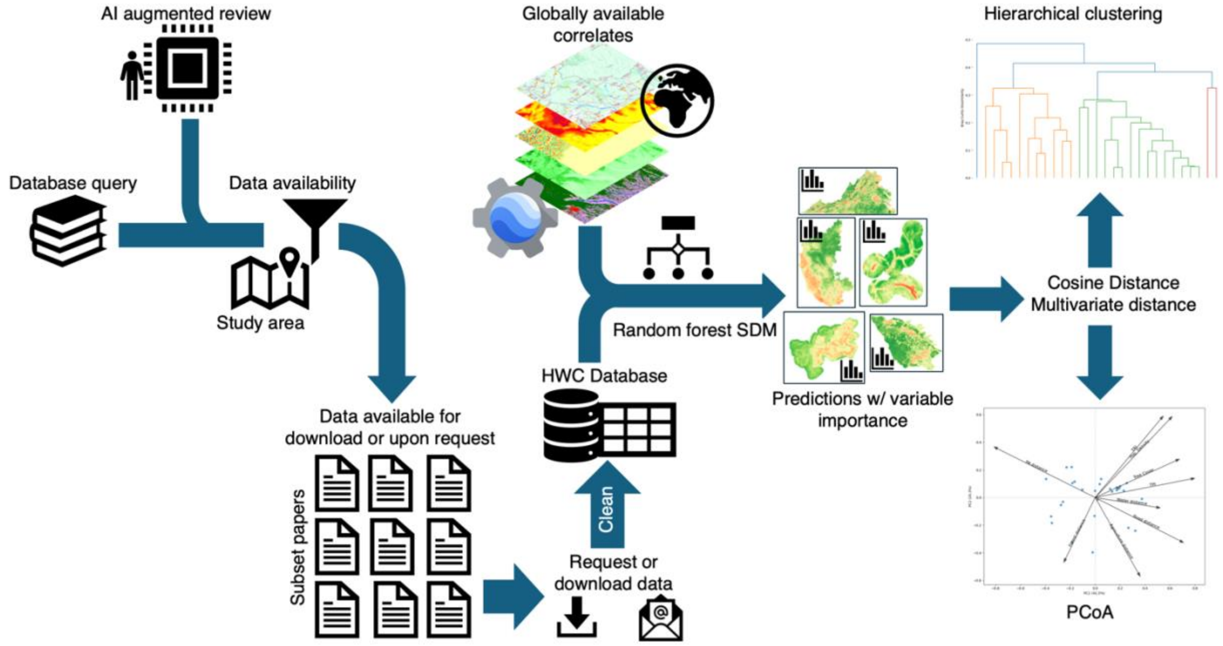


**Figure S2.** Flow chart of methodological process consolidating a human-wildlife conflict database to predicting and interpreting results. Candidate studies were identified through literature review and studies with available data were flagged by an LLM. Data was downloaded and cleaned to be used in a species distribution model for each study area and conflict scenario. Model outputs including variable importance are used to compare conflict scenarios and identify the similarities and differences between scenarios.

### Tables

**Table S1.** Variables considered for analysis – Information including the dates of the data as well as the source and brief descriptions found within the table.

| Variable | Resolution | Years | Data source | Citation |
|---|---|---|---|---|
| Terrain ruggedness index | 30m | 2007 | NASA SRTM + S.J. Riley 1999 | (1) |
| Elevation | 30m | 2007 | NASA SRTM | (2) |
| Multi-scale topographic position index | 270m | 2015 | NASA SRTM derived | (3) |
| Population density | ~926m | 2000-2022 | Gridded population of the world v4 UN WPP adjustment | (4) |
| Human-development index | ~9300m | 1990-2016 | Gridded global GDP per capita v2 | (5) |
| Gross domestic product - Purchasing power parity | ~9300m | 1990-2016 | Gridded global GDP per capita v2 | (5) |
| Livestock density | ~10000m | 2020 | Gridded livestock of the world (GLW4) | (6) |
| Crop productivity | ~10000m | 2010-2020 | Global Gridded Crop Production Dataset | (7) |
| Human modification index | 300m | 1990-2020 | Global extent and change in human modification of terrestrial ecosystems | (8) |
| Median tree canopy cover | 30m | 2000-2015 | Global Forest Cover Change Tree Cover Dataset | (9) |
| Distance from roads | ~10m | 2018 | Global Roads Inventory Project | (10) |
| Distance from forest | 10m | 2021 | ESA WorldCover 2021 | (11) |
| Distance from agriculture | 10m | 2022 | ESA WorldCover 2021 | (11) |
| Distance from shrubland and grassland | 10m | 2023 | ESA WorldCover 2021 | (11) |
| Distance from built-up areas | 10m | 2024 | ESA WorldCover 2021 | (11) |
| Distance from other natural area | 10m | 2025 | ESA WorldCover 2021 | (11) |
| Distance from water | 90m | 2019 | MERIT Hydro | (12) |
| Distance from Protected areas | 10m | 2025 | WDPA IUCN I-IV + Indian PAs | (13) |
| Biome | 1000m | 2018 | Global mapping of potential natural vegetation | (14) |
| Effective mesh size | 1000m | 2021 | ESA WorldCover 2021 -PyLandstats | (11, 15, 16) |
| Edge density | 1000m | 2021 | ESA WorldCover 2021 -PyLandstats | (11, 15, 16) |
| Largest patch size | 1000m | 2021 | ESA WorldCover 2021 -PyLandstats | (11, 15, 16) |
| Patch density | 1000m | 2021 | ESA WorldCover 2021 -PyLandstats | (11, 15, 16) |
| Fractal dimension index | 1000m | 2021 | ESA WorldCover 2021 -PyLandstats | (11, 15, 16) |

**Table S2.** Variance inflation factor results for included variables

| dataset | PA_dist | TCC | agriculture_dist_esa | forest_dist_esa | mTPI | popDens | road_dist | tri | water_dist |
|---|---|---|---|---|---|---|---|---|---|
| **combined** | 1.92 | 1.83 | 1.37 | 1.05 | 1.01 | 1.03 | 1.51 | 2.07 | 1.64 |
| **Athreya_etal_2015** | 1.26 | 2.73 | 2.03 | 1.83 | 1.09 | 1.10 | 1.99 | 2.69 | 2.00 |
| **ConflictValidation_Qnnaire** | 1.71 | 3.88 | 3.12 | 1.58 | 1.09 | 3.21 | 2.00 | 2.57 | 2.53 |
| **coyotes_Farr** | 3.41 | 1.84 | 3.37 | 2.55 | 1.05 | 2.75 | 1.57 | 2.25 | 2.64 |
| **Loveridge_etal_2017_carnivore_pred** | 1.02 | 2.40 | 1.57 | 1.32 | 1.00 | 1.09 | 2.36 | 1.67 | 2.20 |
| **Malviya_Krishnamurthy_2022_tiger** | 1.59 | 4.35 | 2.83 | 1.88 | 1.09 | 6.54 | 2.47 | 2.73 | 3.38 |
| **media_conflict_data_052021** | 1.26 | 2.73 | 2.03 | 1.83 | 1.09 | 1.10 | 1.99 | 2.69 | 2.00 |
| **Api_point_ analysis** | 2.62 | 5.22 | 3.35 | 1.51 | 1.07 | 1.57 | 3.58 | 4.38 | 1.79 |
| **Baral_etal_2021_incidents** | 4.00 | 3.29 | 4.58 | 2.67 | 1.17 | 1.20 | 4.71 | 3.75 | 3.08 |
| **Elephas_maximus_Location** | 2.04 | 5.45 | 3.92 | 1.35 | 1.11 | 1.78 | 3.92 | 2.78 | 2.02 |
| **snow_leopard_ attacks** | 25.40 | 1.53 | 1.29 | 2.33 | 1.43 | 1.73 | 1.98 | 1.14 | 2.14 |
| **Yang_et_al_2024_wildboars** | 11.67 | 1.81 | 1.40 | 1.29 | 1.04 | 1.07 | 1.13 | 1.20 | 1.05 |
| **slothbear_conflict** | 2.55 | 5.66 | 4.20 | 1.42 | 1.12 | 4.26 | 3.32 | 2.26 | 2.69 |
| **Pop_etal_2023a_combo** | 2.83 | 5.75 | 3.26 | 1.47 | 1.02 | 1.23 | 3.44 | 4.41 | 2.30 |
| **Naha_2020_eleph_crops_clean** | 2.94 | 4.65 | 3.05 | 1.42 | 1.02 | 3.32 | 2.10 | 2.65 | 2.16 |
| **CLAWS_Depredation Events_Jun-2016_Jun-2018** | 3.30 | 3.18 | 11.37 | 2.28 | 1.01 | 22.60 | 3.13 | 7.02 | 6.69 |
| **wolf_pred_1999_2009_Treves** | 1.08 | 2.95 | 2.00 | 1.33 | 1.01 | 1.03 | 2.46 | 1.98 | 1.63 |
| **Simon_and_fortin2020_classified** | 10.82 | 11.61 | 12.30 | 2.32 | 1.16 | 2.42 | 18.79 | 3.53 | 3.51 |
| **Davoli_SusScrofa_2022** | 2.50 | 4.31 | 2.55 | 1.58 | 1.06 | 1.08 | 2.47 | 4.54 | 2.55 |
| **Mekonnen_etal_2012_balemonkey_crop** | 4.82 | 4.93 | 1.94 | 1.27 | 1.01 | 3.06 | 2.74 | 3.87 | 2.99 |
| **Branco_etal_2019_elephant** | 2.68 | 4.36 | 2.50 | 1.38 | 1.01 | 5.05 | 2.65 | 2.48 | 2.78 |
| **Puma_kills_Guerisoli2017_reproj** | 0.00 | 1.14 | 1.31 | 1.03 | 1.09 | 1.01 | 1.18 | 1.14 | 1.07 |
| **Wolf_pred_databaseTot** | 2.50 | 4.30 | 2.56 | 1.58 | 1.06 | 1.08 | 2.47 | 4.55 | 2.56 |
| **EC_All** | 2.27 | 5.82 | 2.39 | 1.27 | 1.04 | 1.06 | 2.28 | 3.84 | 1.76 |
